# Comparative Multi-omics Reveals Conserved Membrane Architecture and Selective Metabolic Cargo Enrichment Across Yeast-Derived Extracellular Vesicles

**DOI:** 10.64898/2026.10.23.761364

**Authors:** Mahmoud S. Yahia, Sara P. McLaughlin, Krishay Sachdeva, Sung Hyun Cho, Peter Kozloff, Julia A. Mudryk, Felix Tettey-Wayo, Melanie R. McReynolds

## Abstract

Extracellular vesicles (EVs) are fundamental mediators of intercellular communication across all kingdoms of life, yet our understanding of EV biology has been derived predominantly from mammalian systems. In contrast, yeast-derived extracellular vesicles (YDEVs) remain poorly characterized despite their potential as scalable, non-pathogenic platforms for biotechnology and metabolic engineering. Here, we combined a high-yield isolation workflow with comparative multi-omics to define the metabolically active cargo and membrane composition of EVs derived from three yeast species and four strains: the conventional model yeast *Saccharomyces cerevisiae* (SC), the non-model oleaginous yeast *Rhodotorula toruloides* (RT-88), and two *Yarrowia lipolytica* strains, wild-type W29 and its engineered derivative Po1f. Using tangential flow filtration coupled with ultracentrifugation, we reproducibly isolated intact, bilayer-enclosed vesicles with conserved nanoscale morphology across all strains despite marked differences in EV yield. Comparative metabolomics revealed selective enrichment of distinct metabolically active cargo, including nucleotide metabolites in SC, fatty acids and acyl-carnitines in RT-88, and strain-specific metabolites within the *Yarrowia* isolates. Notably, nicotinamide adenine dinucleotide (NAD⁺) was enriched in SC and RT-88 EVs but was nearly absent from *Yarrowia*-derived vesicles, raising the possibility that EV encapsulation may facilitate protected transport of this otherwise membrane-impermeable cofactor. In contrast to these differences in cargo composition, comparative lipidomics demonstrated a broadly conserved membrane architecture across all strains, with only quantitative remodeling of specific lipid classes and membrane unsaturation profiles. Together, these findings support an organizational principle in which yeast-derived extracellular vesicles preserve a conserved membrane architecture while selectively enriching metabolically active cargo, establishing a comparative framework for understanding EV biology across evolutionarily diverse yeast species.

**Graphical Abstract:** 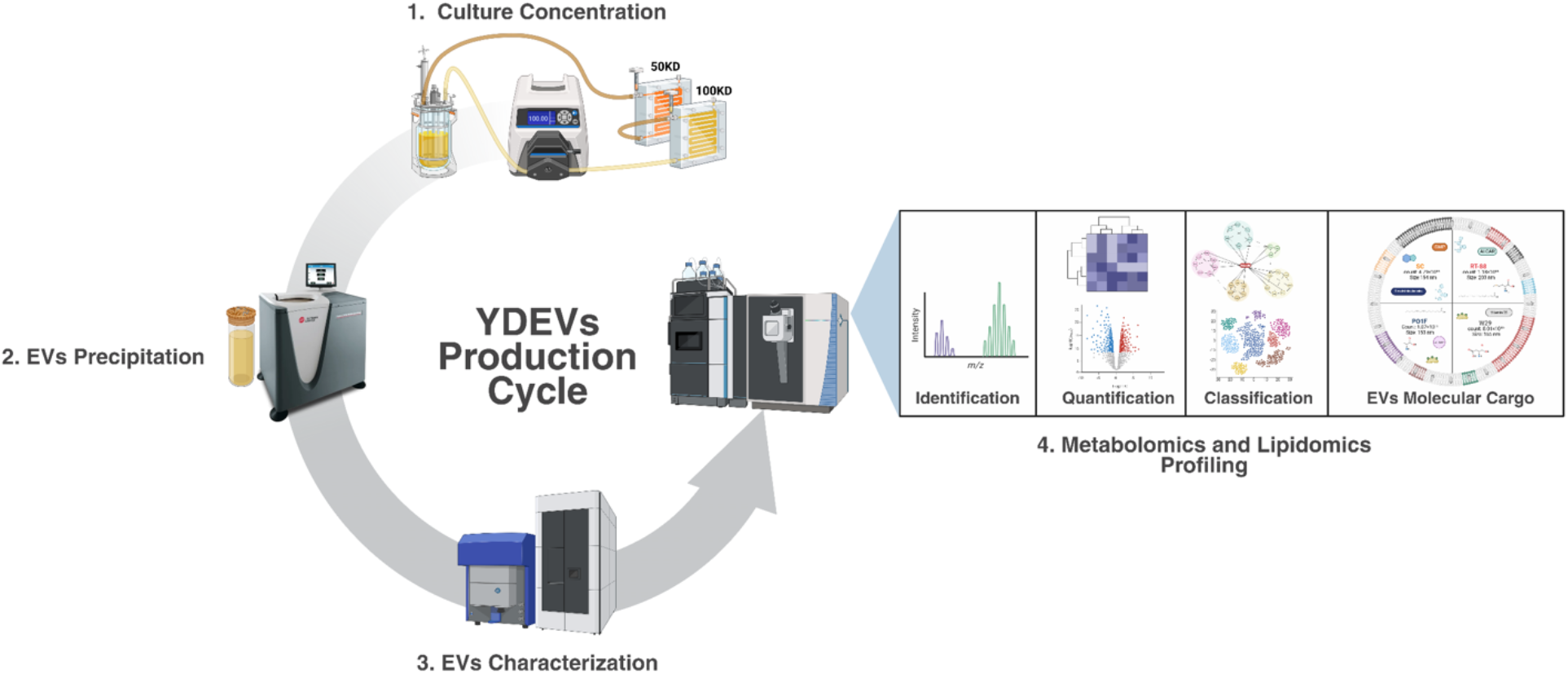

## Introduction

Extracellular vesicles (EVs) are nanoscale, phospholipid bilayer-enclosed particles, typically ranging from 50–300 nm in diameter, that have emerged as fundamental mediators of intercellular communication across all kingdoms of life (Kalluri & LeBleu, 2020). Secreted by virtually all cell types, EVs are generated through evolutionarily conserved biogenesis pathways that, although mechanistically distinct among organisms, converge on a common function: the transfer of biologically active molecular cargo between cells. This cargo—including proteins, nucleic acids, and metabolites, enables EVs to regulate diverse physiological processes by facilitating the exchange of molecular information between cells and their surrounding environment (Doyle & Wang, 2019). Traditionally, EVs have been classified into exosomes, microvesicles, and apoptotic bodies based on their biogenesis, size, and functional heterogeneity. More recently, however, the discovery of non-canonical EV subtypes, including organelle-rich blebbisomes (Jeppesen et al., 2025), metabolically active extracellular “cell couriers” termed zombosomes (Dakhel et al., 2026), and the ultrasmall supermeres (Zhang et al., 2021), has expanded our understanding of EV diversity and highlighted that the biology of extracellular vesicles remains far from fully defined. Importantly, nearly all of our current understanding of EV biology has been derived from mammalian systems.

In contrast, yeast-derived extracellular vesicles (YDEVs) remain comparatively understudied despite their considerable biological and biotechnological potential. Unlike mammalian EVs, YDEVs lack many of the canonical markers routinely used for EV characterization, including the tetraspanins CD9, CD63, and CD81 and the ESCRT (endosomal sorting complexes required for transport)-associated proteins Alix, TSG101, and syntenin (Jankovičová et al., 2020). This distinction is particularly intriguing given that much of our mechanistic understanding of ESCRT-dependent membrane remodeling and exosome biogenesis originates from yeast model systems (Schmidt & Teis, 2012). Despite these foundational discoveries, fungal EVs have not yet been systematically classified into discrete subpopulations (Y. Li et al., 2026). Moreover, although non-pathogenic yeasts represent highly scalable production platforms—with global fermentation capacity exceeding 30 million liters annually compared with approximately 7 million liters of mammalian cell culture capacity (State of Global Fermentation Capacity, 2023), fundamental questions surrounding YDEV biogenesis, cargo selection, and regulatory mechanisms remain unresolved (X. Li et al., 2019). Progress has also been hindered by inconsistent isolation methodologies, limiting reproducibility and making it difficult to compare EV composition across studies or across yeast species (Y. Li et al., 2026).

Addressing these challenges requires expanding EV research beyond mammalian systems to encompass evolutionarily and industrially relevant yeast species. *Saccharomyces cerevisiae*, the conventional model yeast, has emerged as a powerful platform for investigating EV biology because of its genetically tractable genome and extensive use as a microbial cell factory capable of high titer, rate, and yield (TRY) production (Nielsen, 2019). Recent studies have demonstrated that *S. cerevisiae* EV cargo is dynamically remodeled in response to environmental stress, including the enrichment of thermotolerance-associated proteins following heat stress (Logan et al., 2024), while the identification of the SNARE proteins Sso2 and Nyv1 within EV membranes has provided new insight into fungal EV biogenesis and potential biomarkers (Y. Li et al., 2026). Despite these advances, the molecular composition of YDEVs remains largely unexplored. In particular, no study has comprehensively characterized both the metabolically active cargo and membrane composition of EVs across model and industrially relevant yeast species. This gap persists largely because metabolomic and lipidomic analyses require substantially higher EV yields than are typically obtainable from mammalian cell culture or biological fluids, emphasizing the need for scalable production and isolation strategies.

Understanding the biological organization of YDEVs requires examining both the molecular cargo they encapsulate and the membrane architecture that enables its transport. EV function is not dictated by any single molecular component but instead emerges from the coordinated contributions of a heterogeneous cargo packaged within a structurally specialized membrane. Although EV research has historically focused on proteins and nucleic acids, comparatively little is known about how metabolites and membrane lipids are organized within yeast-derived vesicles. Defining these complementary molecular layers is therefore essential for understanding both EV composition and their potential biological function.

Here, we combine a scalable EV isolation platform with comparative multi-omics to characterize YDEVs across three yeast species and four strains: the conventional model yeast *Saccharomyces cerevisiae* (SC), the non-model oleaginous yeast *Rhodotorula toruloides* (RT-88), and two *Yarrowia lipolytica* strains, the wild-type W29 and its engineered derivative Po1f. Using high-throughput tangential flow filtration coupled with ultracentrifugation, we establish a reproducible workflow for high-yield YDEV isolation and apply deep untargeted metabolomic and lipidomic profiling to define the molecular organization of these vesicles. Our analyses reveal that, despite maintaining a broadly conserved membrane architecture, YDEVs exhibit selective enrichment of metabolically active cargo across yeast species and strains. Together, these findings support an organizational principle in which a conserved membrane framework provides a stable structural platform for the selective enrichment of metabolically active cargo across evolutionarily and industrially relevant yeasts.

## Materials and Methods

### Yeast strains and growth kinetics

Four yeast strains were obtained from the Center for Advanced Bioenergy and Bioproducts Innovation (CABBI): the conventional model yeast *Saccharomyces cerevisiae* (SC), the non-model oleaginous yeast *Rhodotorula toruloides* (RT-88), and two *Yarrowia lipolytica* strains, the wild-type W29 and its engineered derivative Po1f. Strains were maintained as glycerol stocks at −80 °C, streaked onto yeast extract peptone dextrose (YPD; 1% yeast extract, 2% peptone, 2% dextrose) agar plates, and incubated at 30 °C for 48 h. Single colonies were inoculated into 5 mL of YPD broth and cultured overnight at 30 °C with orbital shaking at 240 rpm to generate starter cultures for growth-kinetics analysis and extracellular vesicle (EV) production.

Growth kinetics were determined to establish a standardized physiological harvest point for EV isolation. Overnight cultures were diluted in fresh YPD medium to an initial optical density at 600 nm (OD₆₀₀) of 0.05–0.1, and 200 µL aliquots were dispensed into sterile, flat-bottom, non-treated 96-well plates. For each strain, three independent colonies were analyzed with six technical replicates per colony (n = 18 wells per strain). Plates were incubated in a Varioskan LUX Multimode Microplate Reader (Thermo Fisher Scientific, Waltham, MA, USA) at 30 °C with continuous orbital shaking (240 rpm), and optical density at 600 nm (OD₆₀₀) and 650 nm (OD₆₅₀) was recorded every 15 min for 84 h. Wells containing sterile YPD medium served as blanks, and OD₆₀₀ measurements were background corrected by blank subtraction prior to analysis.

Growth curves were analyzed using the Quantitative Growth Curve Evaluation (QurvE v1.1.1) pipeline with nonparametric spline fitting to estimate the maximum specific growth rate (μmax), doubling time, lag phase duration, and area under the growth curve (Wirth et al., 2023). The resulting growth-rate profiles were used to identify a common physiological stage for EV collection across all strains. Based on these analyses, cultures were harvested during the late logarithmic growth phase (approximately 14 h), corresponding to a comparable metabolically active physiological state for downstream EV isolation and characterization (Fig. 1B,C).

**Figure 1.**
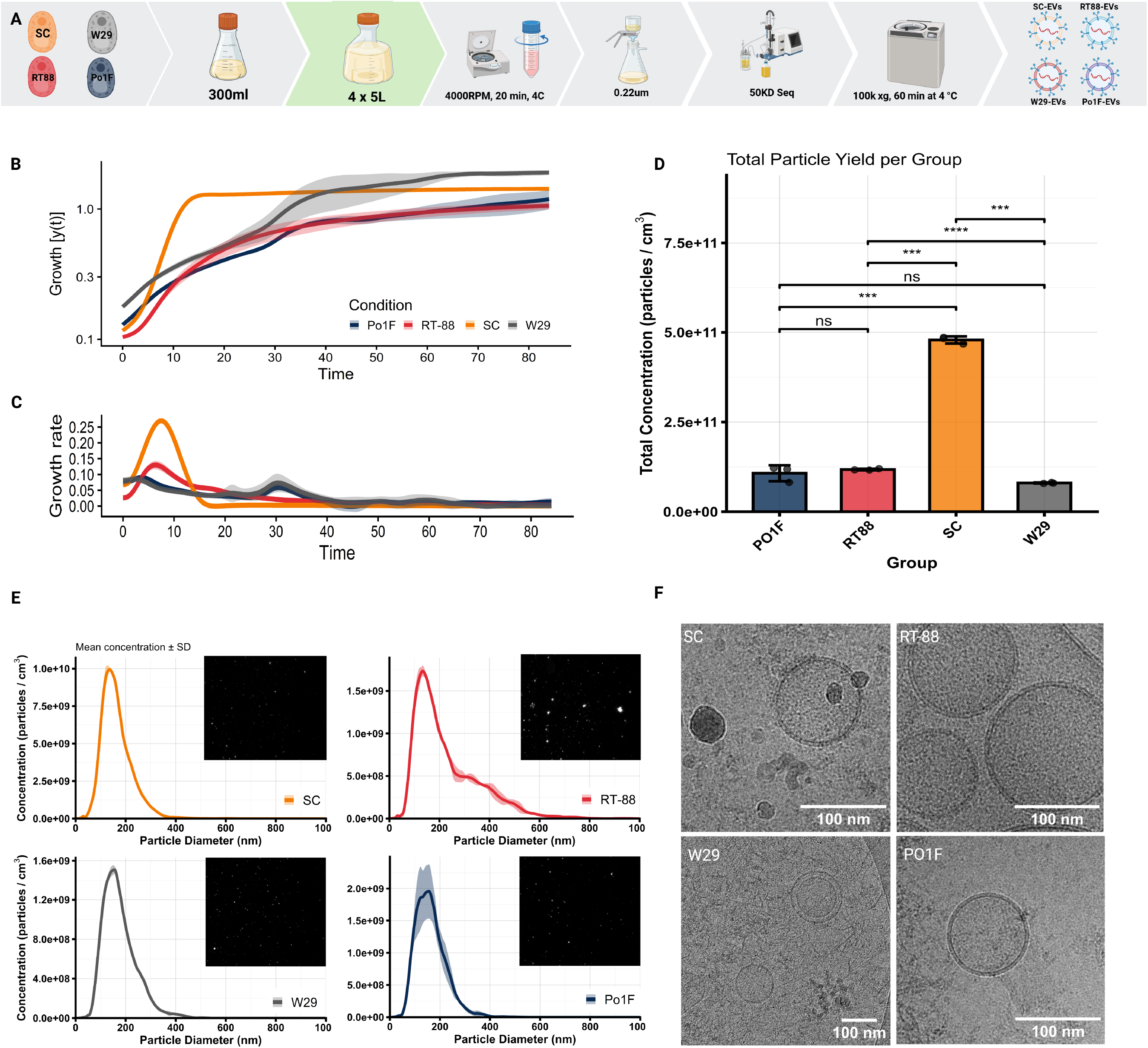
Establishment of a scalable workflow for the isolation and characterization of yeast-derived extracellular vesicles (YDEVs). (A) Schematic of the YDEV isolation workflow, including seed culture expansion, large-scale cultivation (4 × 5 L), clarification, dual-stage tangential flow filtration (100- and 50-kDa molecular weight cutoff), and ultracentrifugation for EV purification. (B, C) Growth kinetics of the four yeast strains used to define a standardized harvest point. Although growth rates differed among strains, cultures were harvested at approximately 14 h, corresponding to a shared late-logarithmic physiological state characterized by converging growth-rate profiles. (D) Nanoparticle tracking analysis (NTA) demonstrating significant differences in total EV production among strains, with *Saccharomyces cerevisiae* (SC) producing approximately four-fold more vesicles than the non-model yeast strains (one-way ANOVA with Tukey’s post hoc test; ns, not significant; \*\**P* < 0.001, \*\*\**P* < 0.0001; n = 3). (E) Representative NTA particle size distributions showing that EVs from all four strains exhibit overlapping nanoscale size distributions with mean particle diameters ranging from 153 to 203 nm. (F) Representative cryo-EM images confirming intact, spherical, membrane-enclosed extracellular vesicles with well-defined phospholipid bilayers across all four yeast strains. Together, these data establish a reproducible and scalable workflow for isolating high-quality YDEVs suitable for comparative multi-omics analyses.

### Isolation and purification of yeast-derived extracellular vesicles (YDEVs)

EVs were isolated from conditioned media collected from all four yeast strains using a scalable tangential flow filtration (TFF) workflow coupled with ultracentrifugation, adapted from previously described methods for concentrating EVs from mammalian cell culture media with modifications (Busatto et al., 2018). To generate production cultures, 1 mL of overnight starter culture was transferred into 250 mL of yeast extract peptone dextrose (YPD) medium and incubated overnight at 30 °C with orbital shaking at 230 rpm. Seed cultures were quantified using an automated cell counter (Countess; Thermo Fisher Scientific) and used to inoculate 2.5 L production cultures in 5 L baffled flasks to a starting OD₆₀₀ of 0.1. Cultures were grown at 30 °C with shaking and harvested during the late logarithmic growth phase, as determined by the growth kinetics analysis.

Cells were removed by centrifugation at 4,000 × *g* for 20 min at 4 °C, and the resulting conditioned medium was clarified through a 0.45 µm Steritop vacuum filtration unit (Millipore) to remove residual cellular debris. Clarified conditioned medium was concentrated using a two-stage TFF system consisting of polyethersulfone (PES) Pierce™ Crossflow cassettes (100 kDa and 50 kDa molecular weight cutoffs) connected in series and operated with a Masterflex peristaltic pump (L/S 16 pump head) at a constant transmembrane pressure of 2 bar. The conditioned medium was continuously recirculated through the system until the EV-containing retentate reached the desired concentration.

The concentrated retentate was subsequently subjected to ultracentrifugation at 100,000 × *g* for 1 hr. at 4 °C to pellet EVs. Pelleted vesicles were resuspended in 1 mL of ice-cold phosphate-buffered saline (PBS), aliquoted into 100 µL fractions, and stored at −80 °C until downstream characterization and multi-omics analyses.

### Extracellular Vesicle Characterization

#### Nanoparticle tracking analysis (NTA)

Particle concentration and size distribution were determined by nanoparticle tracking analysis (NTA) using a ZetaView nanoparticle tracking analyzer and ZetaView software (Particle Metrix, Inning am Ammersee, Germany). Prior to sample acquisition, the instrument was calibrated according to the manufacturer’s recommendations using reference nanoparticle standards, and particle-free phosphate-buffered saline (PBS) was analyzed as a blank to confirm the absence of background signal. EV samples were diluted in PBS to achieve the recommended acquisition range of 140–200 particles per frame. Measurements were acquired using the manufacturer’s default extracellular vesicle settings by scanning 11 camera positions with 60 frames captured per position. Each biological sample was analyzed in technical triplicate, and replicate measurements were averaged for downstream analyses.

#### Cryogenic electron microscopy (cryo-EM)

To assess morphology and membrane integrity, EV samples from all four yeast strains were subjected to cryo-EM Holey carbon grids (Quantifoil, R 1.2/1.3, 300 mesh) were glow-discharged at 15 mA for 50 s immediately before vitrification. A 5 µL aliquot of each EV suspension was applied directly to the grid, incubated for 420 s, blotted for 5 s, and plunge-frozen in liquid ethane with an EM GP2 Plunger (Leica Microsystems) to generate vitrified specimens.

Vitrified grids were imaged on a Titan Krios G3 transmission electron microscope (Thermo Fisher Scientific) operated at 300 kV, equipped with a Falcon 4i direct electron detector and a Selectris X energy filter. Data collection was automated using EPU software (ver. 3.11.0.9330). Micrographs were acquired under low-dose conditions with an exposure rate of 1.05 e^-^/A^2^/s and a total electron dose of 20 e⁻/Å²) at a nominal magnification of 33,000×, yielding a calibrated pixel size of 3.0 Å. A target defocus range of −3.0 to −4.0 µm was applied to maintain membrane integrity while providing sufficient contrast for visualization of intact phospholipid bilayers. Multiple grid squares were examined for each sample to ensure that the observed EV morphology was representative of the entire preparation. Visualization and scaling of the MRC files were performed using the Fiji software package (Schindelin et al., 2012).

### Metabolomics and Lipidomics Analysis by LC-MS/MS

#### Multi-omics sample preparation

Purified EV preparations were aliquoted immediately following isolation and stored at −80 °C until metabolomic and lipidomic analyses. Independent aliquots were used for metabolite and lipid extractions to minimize repeated freeze–thaw cycles. Prior to extraction, EV samples were thawed on ice and processed using optimized workflows for polar metabolite and lipid analyses.

For metabolomic profiling, cold-solvent precipitation method adapted for EV preparations, treated with an ice-cold extraction solvent (40:40:20 methanol:acetonitrile:water supplemented with 0.5% formic acid), and neutralized with 15.8% (w/v) ammonium bicarbonate, vortexed, incubated on dry ice for 10 min, and centrifuged at 16,000 × *g* for 25 min at 4 °C (Lu et al., 2020). The clarified supernatant was collected and analyzed by liquid chromatography–mass spectrometry (LC-MS) within 24 hrs. of extraction.

For lipidomic profiling, EV samples were extracted using isopropanol:H₂O:ethyl acetate (30:10:60, v/v/v) containing Avanti Lipidomix internal standards (Avanti Polar Lipids, Alabaster, AL; 1:1000 dilution) (Liakh et al., 2020). Samples were vortexed, sonicated for 5 min, and centrifuged at 15,000 × *g* for 5 min at 4 °C. The organic phase was collected, and the remaining pellet was subjected to a second extraction using the same procedure. Organic extracts were combined, dried under nitrogen gas, and stored until LC-MS analysis.

#### LC-MS metabolomic profiling

Polar metabolite extracts were analyzed within 24 h using hydrophilic interaction chromatography (HILIC) coupled to an Orbitrap Exploris 240 mass spectrometer (Thermo Fisher Scientific), as previously described (Doyle & Wang, 2019). Chromatographic separation was performed on an XBridge BEH Amide column (2.1 × 150 mm, 3.5 μm; Waters, Milford, MA). Mobile phase A consisted of 95:5 water:acetonitrile containing 20 mM ammonium acetate and 20 mM ammonium hydroxide, whereas mobile phase B consisted of 90:10 acetonitrile:water containing the same additives. Metabolites were separated using the following gradient: 0–2 min, 90% B; 3–7 min, 75% B; 8–9 min, 70% B; 10–12 min, 50% B; 13–14 min, 25% B; 16–20 min, 0% B; and 21–25 min, 90% B for column re-equilibration. The LC system was operated at a flow rate of 150 µL min⁻¹ with a column temperature of 25 °C, an autosampler temperature of 5 °C, and an injection volume of 5 µL. Mass spectrometry data were acquired in both positive and negative electrospray ionization modes at a resolution of 180,000 (at *m/z* 200), an automatic gain control (AGC) target of 3 × 10⁶, a maximum injection time of 30 ms, and a scan range of *m/z* 70–1000.

#### LC-MS lipidomic profiling

Dried lipid extracts were reconstituted in 300 µL of solvent A (isopropanol:acetonitrile:H₂O, 45:35:20, v/v/v), vortexed, sonicated for 7 min, and centrifuged twice at 15,000 × *g* to remove insoluble material. Aliquots (60 µL) of clarified extracts were transferred to LC-MS vials for analysis. Lipidomic analyses were performed within 36 hrs. of extraction using a Vanquish UHPLC system coupled to an Orbitrap Exploris 240 mass spectrometer equipped with an H-ESI ion source (Thermo Fisher Scientific). Lipids were separated on a Waters CSH C18 column (1.0 × 150 mm, 1.7 μm particle size). Mobile phase A consisted of acetonitrile:H₂O (60:40, v/v) containing 10 mM ammonium formate and 0.1% formic acid, whereas mobile phase B consisted of isopropanol:acetonitrile (95:5, v/v) containing 10 mM ammonium formate and 0.1% formic acid. The mobile phase was delivered at 0.11 mL min⁻¹ with the column maintained at 65 °C. The solvent B gradient was programmed as follows: 15% (0 min), 30% (2 min), 48% (2.5 min), 82% (11 min), 99% (11.01–12.95 min), and returned to 15% for column equilibration (13–15 min). Mass spectrometry data were acquired in both positive and negative ionization modes using spray voltages of 4.0 kV and 3.0 kV, respectively. Full-scan spectra were acquired over an *m/z* range of 200–1000, and data-dependent MS/MS spectra were collected using AcquireX with a stepped collision energy of 30% (±5%) to facilitate lipid annotation.

### Data Processing and Statistical Analysis

#### Spectral processing and compound annotation

Raw UHPLC-MS/MS data from metabolomic and lipidomic analyses were processed using MS-DIAL (v5.5.260323) for feature detection, chromatographic alignment, peak integration, and compound annotation (Tsugawa et al., 2015). Features were detected using a minimum peak-height threshold of 1 × 10⁵, and blank subtraction together with ghost-peak removal were applied to minimize background and artifactual signals. Metabolites were annotated using the comprehensive MS-DIAL metabolite spectral library, whereas lipid species were annotated using the integrated MS-DIAL lipid database. For lipidomic analyses, EquiSPLASH™ internal standards were incorporated during feature processing to normalize lipid signal intensities for quantitative comparisons. All metabolite and lipid annotations were manually inspected and curated prior to export of aligned feature tables for downstream analyses. Metabolomic datasets included features detected across both positive and negative ionization modes, whereas lipidomic analyses were performed using positive-ion acquisition.

#### Metabolomic data processing

Metabolomic feature tables were processed using a custom modular workflow implemented in R (v4.5.3) (R Core Team, 2026). Positive- and negative-ion datasets were merged, and redundant features detected in both ionization modes were removed by retaining the higher-intensity annotation, yielding a curated matrix of 254 metabolite features across 20 biological samples (n = 5 per strain). Features were retained if detected in at least 50% of samples within one biological group. Remaining missing values were imputed using quantile regression imputation of left-censored data (QRILC) (Wei et al., 2018a). To account for differences in EV input, metabolite abundances were normalized to total EV protein determined by bicinchoninic acid (BCA) assay, followed by probabilistic quotient normalization (PQN) (Dieterle et al., 2006). Normalized data were subsequently log₁₀-transformed and autoscaled prior to statistical analyses. Metabolite pathway associations were evaluated using two complementary approaches applied to the normalized (BCA- and PQN-corrected), non-autoscaled feature matrix. Metabolite names were mapped to KEGG compound identifiers by exact-synonym matching against the KEGG REST API (Kanehisa & Goto, 2000). Because the non-conventional strains (RT-88, W29, and Po1f) lack complete species-specific pathway databases, the *Saccharomyces cerevisiae* (sce) KEGG metabolism library was used as a common reference framework for all four strains. In the first approach, rank-based one-versus-rest metabolite set enrichment analysis (MSEA) was performed for each strain by ranking all metabolites according to the log₂ fold change of the strain mean relative to the mean of the remaining three strains, followed by rank-based enrichment testing (fgsea; minSize = 3, maxSize = 500) (Xia & Wishart, 2010). In the second approach, multi-group quantitative enrichment analysis (QEA) was performed using a GlobalTest model (globaltest R package) evaluated across the four strain groups for each pathway containing at least three measured members (Xia et al., 2009).

#### Lipidomic data processing

Lipidomic feature tables were processed using an analogous workflow in R. Because lipid abundances were normalized to EquiSPLASH™ internal standards during MS-DIAL processing, internal-standard features were removed prior to downstream analyses, and no additional normalization to EV protein or PQN was performed to avoid redundant normalization. Following feature de-duplication, 487 lipid species were retained across 20 biological samples. Lipids detected in at least 50% of samples within one biological group were retained for analysis, and remaining missing values were imputed using QRILC. Data were subsequently log₁₀- transformed and autoscaled prior to statistical analyses (Wei et al., 2018b).

#### Statistical analysis

All statistical analyses were performed in R unless otherwise indicated. Differential abundance analyses were designed to identify metabolites and lipid species selectively enriched within extracellular vesicles isolated from each yeast strain, thereby defining strain-associated metabolic cargo while preserving a comparative framework across all four yeast-derived EV populations. Unsupervised principal component analysis (PCA) was used to evaluate global variation in metabolomic and lipidomic profiles across the four yeast strains. Differential abundance was assessed using one-way analysis of variance (ANOVA) followed by Tukey’s honestly significant difference (HSD) post hoc test for pairwise strain comparisons. Although Shapiro–Wilk testing indicated departures from normality for some features, ANOVA was performed on log₁₀-transformed and autoscaled datasets, as the method is robust to moderate deviations from normality in balanced experimental designs. *P* values were adjusted for multiple testing using the Benjamini–Hochberg false discovery rate (FDR) procedure. Features with an FDR-adjusted *P* < 0.05 and an absolute fold change ≥ 1.5 (log₂ fold change calculated from group mean intensities) were considered significantly differentially abundant.

Metabolite pathway enrichment analysis was performed using MetaboAnalyst 6.0 with the Kyoto Encyclopedia of Genes and Genomes (KEGG) pathway database, whereas lipid set enrichment analysis was conducted using the Lipid Set Enrichment Analysis (LSEA) module within LipSig (Lin et al., 2021).

### Data Availability

The raw LC-MS/MS metabolomics and lipidomics datasets generated during this study have been deposited in the MassIVE repository under accession number MSV000102366 and are publicly available upon publication. Processed metabolite and lipid feature tables used for downstream statistical analyses are provided as Supplementary Data.

All custom R scripts and computational workflows used for metabolite and lipid feature curation, polarity integration, feature de-duplication, normalization, statistical analyses, and pathway enrichment analyses have been deposited in a publicly accessible GitHub repository and archived through Zenodo (DOI: 10.5281/zenodo.21199436) to ensure long-term accessibility and computational reproducibility.

## Results

### YDEVs isolation and characterization

To enable comparative multi-omics analyses across yeast species, we first established a reproducible and scalable workflow for the isolation of yeast-derived extracellular vesicles (YDEVs) using tangential flow filtration coupled with ultracentrifugation (Fig. 1A). Because extracellular vesicle composition is influenced by cellular physiology, we next sought to standardize vesicle collection across strains by harvesting cultures at a comparable physiological state rather than at an identical growth rate. Growth kinetics were therefore monitored over approximately 84 hrs. for each yeast strain (Fig. 1B). Although the strains exhibited distinct growth trajectories, their growth-rate profiles converged during the late logarithmic phase at approximately 14 h, representing a metabolically active stage common to all four strains (Fig. 1C). All subsequent EV isolations were performed at this standardized harvest point.

We next assessed EV production and physical characteristics by nanoparticle tracking analysis (NTA). Despite identical inoculation densities, culture volumes, and harvest conditions, EV secretion differed significantly among the four yeast strains (Fig. 1D). *Saccharomyces cerevisiae* (SC) produced the greatest particle yield (4.79 × 10¹¹ particles/cm³), approximately four-fold greater than either *Rhodotorula toruloides* (RT-88; 1.18 × 10¹¹ particles/cm³) or *Yarrowia lipolytica* Po1f (1.07 × 10¹¹ particles/cm³), while the wild-type *Y. lipolytica* W29 strain produced the lowest particle concentration (8.01 × 10¹⁰ particles/cm³). In contrast to these differences in secretion efficiency, EV size distributions were broadly conserved across strains (Fig. 1E). Mean particle diameters ranged from 153–165 nm for SC, W29, and Po1f, whereas RT-88 exhibited a modestly broader distribution with a mean diameter of 203 nm. Importantly, all preparations fell within the expected size range for extracellular vesicles.

To further assess vesicle integrity, cryo-EM was performed on EV preparations from each strain. Representative micrographs confirmed the presence of intact, spherical, membrane-enclosed vesicles with well-defined phospholipid bilayers across all four yeast strains (Fig. 1F). Together, these results establish a robust and scalable workflow for the reproducible isolation of high-quality YDEVs, providing a standardized platform for defining how metabolically active cargo and membrane architecture are organized across evolutionary and industrially relevant yeast species

### Comparative metabolomics reveals selective enrichment of metabolically active cargo across YDEVs

To determine whether YDEVs exhibit conserved or distinct metabolic cargo profiles, we performed untargeted high-resolution LC-MS/MS metabolomics, identifying 254 high-confidence metabolite features across the four yeast strains. Unsupervised principal component analysis (PCA) resolved four well-separated clusters with tight within-group reproducibility (PC1 = 41.6%, PC2 = 32.1%; Fig. 2C), demonstrating that each yeast strain possesses a distinct EV metabolomic profile despite exhibiting broadly similar physical characteristics (Fig. 1, S1). Hierarchical clustering further supported this separation, revealing two higher-order groupings in which the two *Yarrowia lipolytica* strains (W29 and Po1f) clustered together, while *Saccharomyces cerevisiae* (SC) and *Rhodotorula toruloides*(RT-88) formed a second cluster (Fig. 2A). Consistent with these relationships, metabolite ontology analysis demonstrated distinct shifts in metabolite class composition across strains, although a substantial proportion of detected features remained unannotated (Fig. 2D).

**Figure 2.**
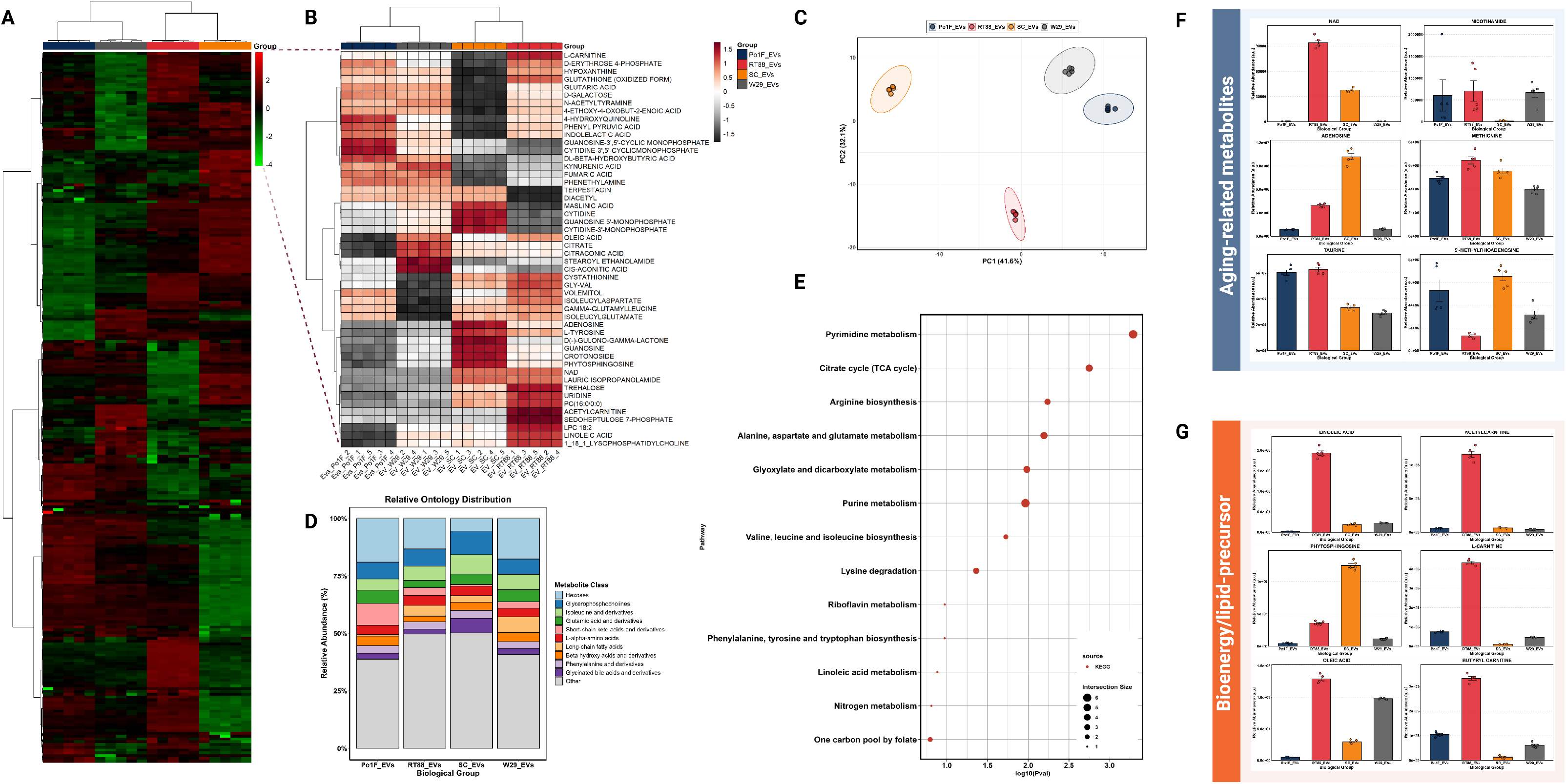
Comparative metabolomics reveals selective enrichment of metabolically active cargo across yeast-derived extracellular vesicles. **(A)** Hierarchical clustering heatmap of 254 annotated metabolite features demonstrating distinct metabolic cargo profiles across extracellular vesicles isolated from the four yeast strains. **(B)** Heatmap of the 50 most differentially abundant metabolites highlighting strain-associated enrichment of metabolically active cargo, including nucleotide metabolites, redox cofactors, amino acids, and lipid-associated metabolites. **(C)** Principal component analysis (PCA) of normalized metabolomic data showing clear separation of EV populations according to yeast strain, indicating distinct metabolic cargo composition across the four YDEV populations. **(D)** Relative metabolite ontology distribution illustrating differences in the chemical composition of EV-associated metabolites while highlighting the diversity of metabolite classes represented within each strain. **(E)** Metabolite over-representation analysis (ORA) identifying significant enrichment of pathways associated with central carbon metabolism, nucleotide metabolism, amino acid metabolism, and cellular redox processes. **(F)** Relative abundance of selected metabolites illustrating strain-associated enrichment of nucleotide metabolites, cofactors, and central metabolic intermediates, including NAD⁺. **(G)** Relative abundance of lipid-associated metabolites demonstrating enrichment of free fatty acids, lysophospholipids, and acyl-carnitines, particularly within *Rhodotorula toruloides* EVs. Together, these analyses demonstrate that YDEVs exhibit distinct, strain-associated metabolic cargo profiles characterized by the selective enrichment of metabolically active metabolites.

We next sought to identify metabolites selectively enriched within YDEVs from each yeast strain. Examination of the 50 most differentially abundant metabolites revealed that EVs were enriched in diverse metabolically active cargo rather than metabolites typically associated with cellular waste (Fig. 2B). Among the most prominent features, nicotinamide adenine dinucleotide (NAD⁺) was selectively enriched in SC and RT-88 EVs but was nearly absent from both *Yarrowia* strains (Fig. 2F). Given that NAD⁺ is generally considered membrane impermeable, its enrichment within membrane-bound extracellular vesicles raises the possibility that EV encapsulation may provide a mechanism for protected extracellular transport of this central metabolic cofactor.

Beyond NAD⁺, each yeast species exhibited selective enrichment of distinct metabolite classes. SC EVs were enriched in purine and pyrimidine metabolites, including adenosine, guanosine, cytidine, and their corresponding monophosphates, together with phytosphingosine and 5′- methylthioadenosine (Fig. 2B,F). SC EVs also contained elevated levels of pentose phosphate pathway intermediates, including sedoheptulose-7-phosphate and D-erythrose-4-phosphate, consistent with enrichment of metabolites associated with nucleotide biosynthesis and central carbon metabolism.

In contrast, EVs isolated from the non-model oleaginous yeast RT-88 exhibited selective enrichment of lipid-associated metabolites, including the free fatty acids oleic acid and linoleic acid, lysophosphatidylcholines, and multiple acyl-carnitines, including acetylcarnitine, L- carnitine, and butyrylcarnitine (Fig. 2G). RT-88 and SC EVs additionally shared enrichment of oxidized glutathione, trehalose, and other metabolites associated with redox homeostasis, whereas taurine was most abundant in RT-88 and Po1f EVs (Fig. 2B). Collectively, these findings indicate that YDEVs selectively enrich metabolically active cargo that reflects the metabolic characteristics of each yeast strain while maintaining a common extracellular vesicle platform.

To place these strain-associated metabolic signatures into a broader biological context, we performed metabolite over-representation analysis (ORA). Significantly enriched pathways included pyrimidine metabolism, the citrate (TCA) cycle, arginine biosynthesis, alanine, aspartate and glutamate metabolism, purine metabolism, and one-carbon metabolism (Fig. 2E). Rather than representing isolated metabolite differences, these pathway-level enrichments indicate that YDEV cargo is preferentially enriched in metabolites associated with central carbon metabolism, nucleotide biosynthesis, amino acid metabolism, and cellular redox homeostasis.

Together, these findings demonstrate that, although YDEVs share conserved extracellular vesicle morphology (Fig. 1), they exhibit selective enrichment of metabolically active cargo across yeast species, establishing metabolic composition as a major distinguishing feature of yeast-derived extracellular vesicles.

### Comparative lipidomics reveals a conserved membrane architecture with quantitative remodeling across YDEVs

Having established that YDEVs exhibit selective enrichment of metabolically active cargo (Fig. 2), we next asked whether this biochemical diversity extends to the membrane itself. Untargeted high-resolution lipidomic profiling identified 487 lipid species across the four yeast strains. In contrast to the pronounced differences observed in metabolite composition, principal component analysis (PCA) revealed substantially greater similarity among lipidomes (Fig. 3B) with higher positive correlation between the samples and the features (Fig. S2), suggesting that YDEVs maintain a broadly conserved membrane architecture despite differences in their internal cargo.

**Figure 3.**
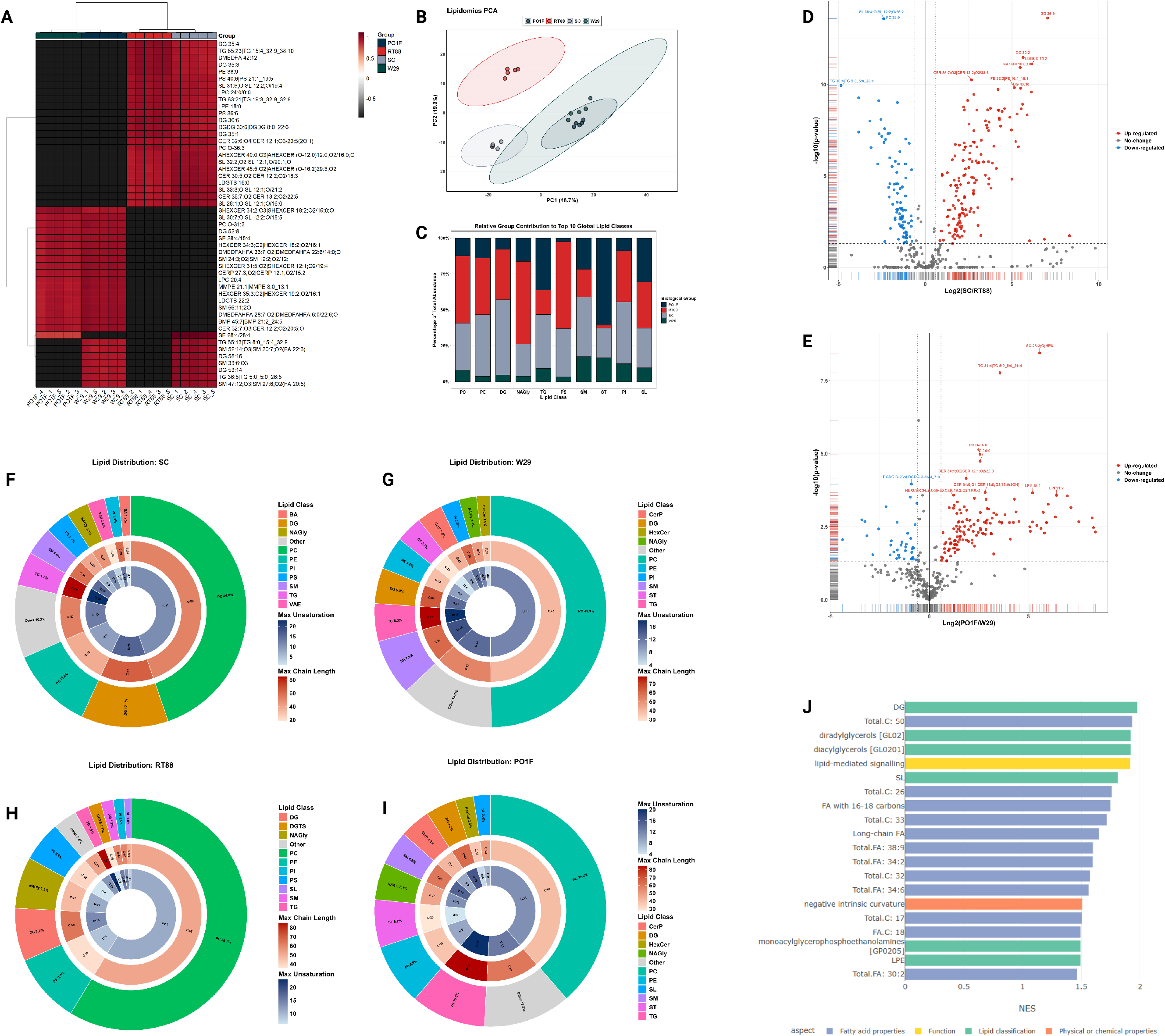
Comparative lipidomics reveals a conserved membrane architecture with quantitative remodeling across yeast-derived extracellular vesicles. **(A)** Heatmap of the 50 most differentially abundant lipid species illustrating quantitative differences in membrane lipid composition across the four yeast strains. **(B)** Principal component analysis (PCA) of normalized lipidomic data demonstrating that YDEV membrane lipidomes are more highly conserved than metabolomic profiles, with the two *Yarrowia lipolytica* strains clustering together and *Saccharomyces cerevisiae* (SC) clustering with *Rhodotorula toruloides*(RT-88). **(C)** Relative lipid class distribution showing that all YDEV populations share a common membrane framework dominated by phosphatidylcholine, phosphatidylethanolamines, sphingolipids, and neutral lipids, while differing in the relative abundance of individual lipid classes. **(D, E)** Pairwise differential abundance analyses highlighting quantitative remodeling of conserved lipid species between SC and RT-88 and between W29 and Po1f, respectively. **(F–I)** Acyl-chain composition analyses demonstrating highly conserved chain-length distributions across strains together with quantitative differences in membrane lipid unsaturation, indicative of strain-associated tuning of membrane fluidity. **(J)** Lipid Set Enrichment Analysis (LSEA) identifying significant enrichment of glycerolipid, glycerophospholipid, sphingolipid, and glycosylphosphatidylinositol (GPI)-anchor biosynthetic pathways. Together, these analyses demonstrate that YDEVs preserve a broadly conserved membrane architecture while quantitatively remodeling lipid composition across evolutionarily and industrially relevant yeast species.

Consistent with this observation, the two *Yarrowia lipolytica* strains (W29 and Po1f) clustered closely, indicating highly similar membrane lipid compositions despite the distinct metabolomic signatures observed in Figure 2. Likewise, *Saccharomyces cerevisiae* (SC) and the oleaginous yeast *Rhodotorula toruloides* (RT-88) formed a second closely related cluster, indicating that membrane lipid composition remains comparatively conserved even among evolutionarily and metabolically distinct yeast species.

To identify the lipid species contributing to this variation, we examined the 50 most differentially abundant lipids across all strains (Fig. 3A). The principal sources of variation included neutral storage lipids, membrane phospholipids, and sphingolipids, including diglycerides (DG), triglycerides (TG), phosphatidylserines (PS), lysophosphatidylcholines (LPC), ceramides, hexosylceramides, and sphingomyelins (Fig. 3A). These observations indicate that quantitative differences among YDEVs arise primarily through remodeling of shared lipid classes rather than the emergence of strain-specific membrane components.

This conclusion was further supported by lipid class analysis (Fig. 3C,F–I). All four strains shared a common membrane framework dominated by phosphatidylcholine (PC), together with phosphatidylethanolamines (PE), sphingolipids, and neutral lipids. Although the relative abundance of individual lipid classes differed among strains—for example, SC EVs were relatively enriched in diglycerides, whereas W29 and Po1f contained proportionally greater sterol and sphingomyelin content—the overall membrane architecture remained highly conserved. These findings suggest that YDEVs assemble their membranes from a common repertoire of structural lipids while quantitatively remodeling individual lipid classes across yeast strains.

To further define this remodeling, we performed pairwise differential abundance analyses between the major PCA groupings (Fig. 3D,E). Comparison of SC and RT-88 identified 150 significantly enriched and 89 depleted lipid species, with RT-88 exhibiting greater abundance of neutral storage lipids, including triglycerides and diacylglyceryl-N,N,N-trimethylhomoserine (DGTS) species. Similarly, comparison of the parental W29 and engineered Po1f strains identified 128 enriched and 37 depleted lipid species, including reduced abundance of several phosphatidylinositol (PI) species in Po1f relative to W29. Together, these findings indicate that strain-associated differences arise primarily through quantitative remodeling of conserved lipid classes rather than wholesale changes in membrane composition.

We next examined membrane biophysical properties by comparing acyl-chain length and unsaturation across strains (Fig. 3F–I). Acyl-chain length was remarkably conserved, with medium-chain lipids (31–40 total carbons) comprising the majority of lipid species in every strain and abundance-weighted mean chain lengths ranging from 33.9 to 35.8 carbons. In contrast, membrane unsaturation varied quantitatively across strains. SC exhibited the most saturated membrane profile, whereas RT-88 contained the greatest proportion of polyunsaturated lipids, with the two *Yarrowia* strains displaying intermediate levels of unsaturation. These findings suggest that YDEVs preserve a conserved membrane framework while modulating membrane fluidity through quantitative changes in lipid unsaturation.

Finally, lipid set enrichment analysis identified significant enrichment of diacylglycerols and related glycerolipid pathways together with lipid-mediated signaling categories (Fig. 3J). Collectively, these results demonstrate that YDEVs maintain a conserved membrane architecture composed of a shared repertoire of structural lipids while selectively remodeling membrane composition through quantitative differences in lipid abundance and unsaturation. When considered alongside the metabolomic analyses (Fig. 2), these findings support an organizational principle in which selective enrichment of metabolically active cargo occurs within a broadly conserved extracellular vesicle membrane framework.

### Comparative multi-omics supports an organizational principle for yeast-derived extracellular vesicles

Collectively, these findings support a fundamental organizational principle in which yeast-derived extracellular vesicles preserve a broadly conserved membrane architecture while selectively enriching metabolically active cargo across evolutionarily and industrially relevant yeast species (Fig. 4). To compare the functional impact of this strain-specific cargo, we mapped the metabolites distinguishing each strain onto their associated *S. cerevisiae* KEGG pathways as a common reference framework, since the non-conventional strains RT-88, W29, and Po1f lack complete species-specific pathway databases. Although all four strains produce nanoscale vesicles with comparable physical characteristics and a shared membrane framework, their molecular cargo exhibited distinct patterns of metabolite enrichment that mapped onto strain-specific pathway signatures. EVs from the model strain SC were distinguished by pyrimidine metabolism, one-carbon folate metabolism, and cofactor biosynthesis, consistent with their nucleoside and cofactor cargo (adenosine, GMP, NAD⁺, 5′-methylthioadenosine). The oleaginous *R. toruloides* (RT-88) ranked highest for glutathione metabolism, the pentose phosphate pathway, vitamin B6 metabolism, and aromatic amino-acid and cysteine/methionine biosynthesis, mirroring its free-fatty-acid, acyl-carnitine, and methionine cargo. The two *Y. lipolytica* strains diverged both from the other strains and from each other: wild-type W29 was defined by central-carbon and organic-acid pathways (TCA cycle, porphyrin metabolism, and glyoxylate/dicarboxylate metabolism), whereas engineered Po1f was distinguished by amino-acid-catabolic pathways (lysine degradation, butanoate, phenylalanine, and sulfur metabolism), alongside taurine and cyclic-nucleotide cargo. Together, this comparative multi-omics framework establishes both a foundational resource for yeast extracellular vesicle biology and a conceptual model in which a conserved membrane architecture provides a stable structural platform for the selective enrichment of metabolically active cargo.

**Figure 4.**
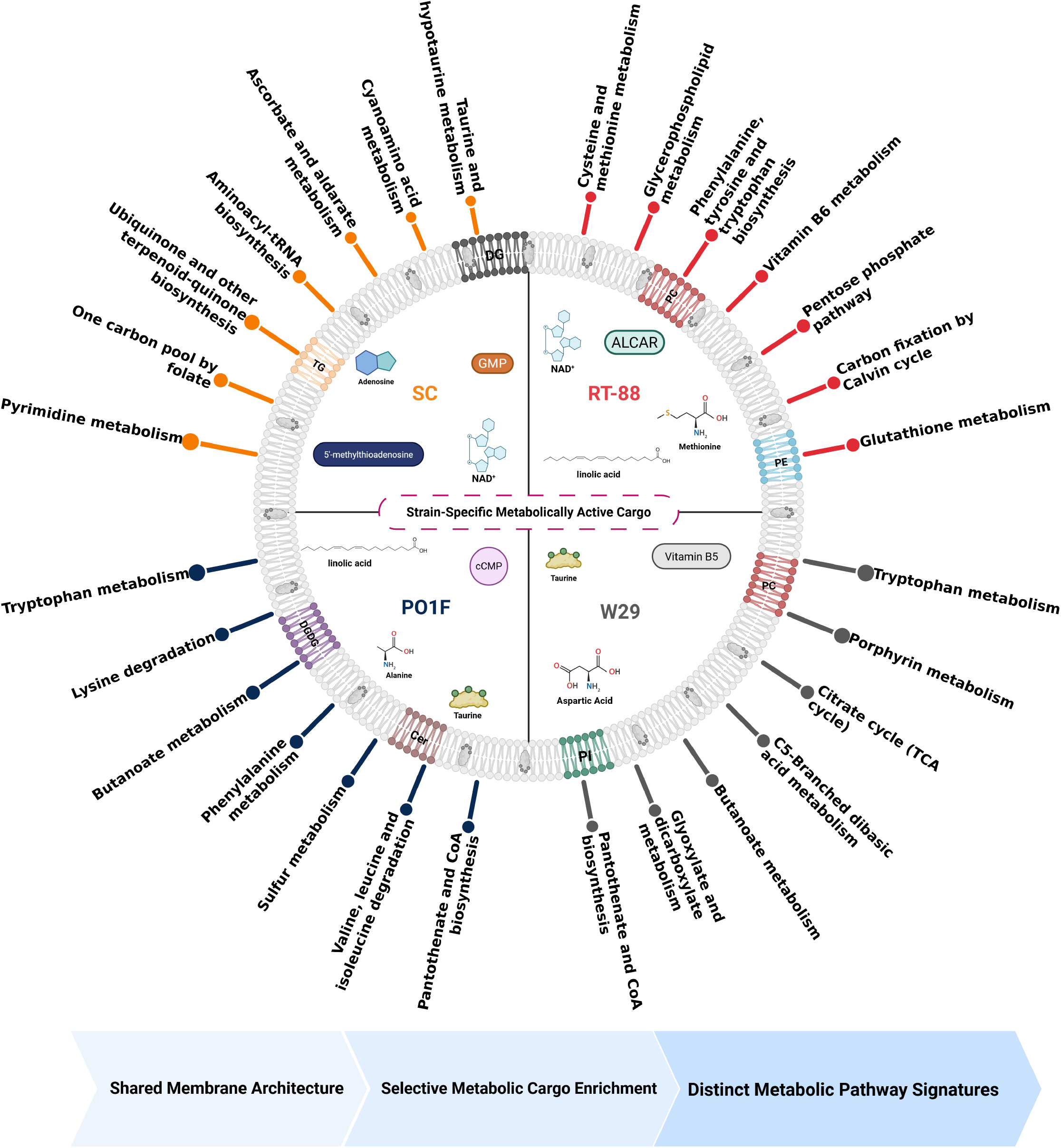
An organizational principle for yeast-derived extracellular vesicles. Yeast-derived extracellular vesicles share a conserved vesicle framework but carry strain-specific molecular cargo. Each quadrant represents the extracellular vesicles of a separate strain (S. cerevisiae (SC), R. toruloides (RT-88), and the two Yarrowia lipolytica strains W29 and Po1f) shown together within a single vesicle schematic to emphasize the conserved bilayer architecture common to all four. Representative enriched metabolites and membrane lipid classes are drawn in each quadrant and colour-coded by class, illustrating that while the vesicle framework is conserved, each strain packages a distinct metabolic and lipidomic payload that can affect the spiked pathways above each quadrant.

## Discussion

The comparative multi-omics analyses presented here support a model in which yeast-derived extracellular vesicles preserve a conserved membrane architecture while selectively enriching metabolically active cargo across evolutionarily and industrially relevant yeast species. This organizational principle provides new insight into the molecular design of fungal extracellular vesicles and establishes a framework for understanding how a conserved membrane platform can support diverse metabolic functions across distinct yeast lineages. Rather than functioning as structurally distinct vesicle populations, our findings suggest that YDEVs are assembled from a common membrane framework that is quantitatively remodeled while supporting selective enrichment of strain-associated metabolic cargo.

One of the most notable findings of this study is that metabolic diversity among YDEVs arises primarily through selective enrichment of metabolically active cargo rather than wholesale changes in vesicle structure. Although all four yeast strains produced morphologically similar extracellular vesicles, each EV population exhibited a distinct metabolite profile enriched for molecules consistent with the metabolic characteristics of the producing strain. EVs from the model yeast *Saccharomyces cerevisiae* preferentially contained nucleotide intermediates and metabolic cofactors, whereas EVs from the oleaginous yeast *Rhodotorula toruloides* were enriched in fatty acids, acyl-carnitines, and other lipid-associated metabolites. Together, these findings support the emerging view that extracellular vesicles function as selective carriers of biologically relevant metabolites rather than passive repositories of cellular by-products.

Among the metabolites identified, the selective enrichment of nicotinamide adenine dinucleotide (NAD⁺) within SC and RT-88 EVs was particularly intriguing. Because NAD⁺ is generally considered membrane impermeable, encapsulation within extracellular vesicles raises the possibility that vesicle-mediated transport could facilitate extracellular distribution of this central metabolic cofactor. Although the functional consequences of EV-associated NAD⁺ remain unknown, these observations provide a rationale for future studies investigating whether YDEVs participate in metabolic communication through cofactor transfer and whether EV-mediated metabolite delivery contributes to cellular adaptation during physiological stress.

In contrast to the pronounced differences observed in metabolite composition, lipidomic analyses revealed remarkable conservation of membrane organization across all four yeast strains. Every YDEV population shared a common membrane framework composed of similar lipid classes and highly conserved acyl-chain length distributions, while quantitative remodeling of lipid abundance and membrane unsaturation represented the principal sources of variation. These observations suggest that evolutionary pressures may preferentially preserve the structural features required for extracellular vesicle stability while allowing sufficient flexibility in membrane composition to accommodate diverse metabolically active cargo. Rather than constructing fundamentally different vesicle membranes, each strain appears to quantitatively remodel a conserved lipid scaffold.

This apparent separation between membrane architecture and cargo composition has important implications for extracellular vesicle engineering. The comparison between the parental *Yarrowia lipolytica* W29 strain and its engineered derivative Po1f demonstrates that substantial remodeling of metabolic cargo can occur while preserving the overall membrane framework and nanoscale morphology of the vesicle. Likewise, the enrichment of fatty acids, acyl-carnitines, and neutral lipids within RT-88 EVs highlights the potential of oleaginous yeasts as scalable production platforms for extracellular vesicles enriched in industrially relevant metabolic intermediates. Together, these findings suggest that YDEVs may function as modular biological delivery platforms whose molecular cargo can be tuned through strain selection or genetic engineering while maintaining a conserved structural chassis.

The scalable isolation workflow developed in this study further expands opportunities for investigating fungal extracellular vesicles beyond traditional model organisms. Although previous studies have largely focused on protein cargo and pathogenic fungi, the integration of high-yield tangential flow filtration with quantitative metabolomics and lipidomics enables systematic characterization of EV composition across diverse yeast species. This analytical framework should facilitate future comparative studies aimed at understanding how metabolism, genetic engineering, and environmental perturbations influence extracellular vesicle composition and function.

Several limitations should be considered when interpreting these findings. First, this study defines the molecular composition of YDEVs but does not directly evaluate vesicle uptake or functional activity in recipient cells. Consequently, the biological significance of the identified metabolites and lipids remains to be established experimentally. Second, although the scalable isolation workflow enabled comparative analyses across multiple yeast species, optimization of upstream cultivation conditions will likely further improve vesicle production from lower-yield strains and enhance their utility for industrial applications. Finally, because lipidomic analyses were performed primarily in positive-ion mode, additional characterization of negatively charged phospholipids and other low-abundance lipid classes may further refine the membrane landscape presented here.

Collectively, this work establishes a comparative molecular framework for yeast-derived extracellular vesicles spanning both model and industrially relevant yeast species. By integrating scalable EV production with quantitative metabolomics and lipidomics, we demonstrate that YDEVs preserve a conserved membrane architecture while selectively enriching metabolically active cargo. These findings establish a foundation for future investigations into extracellular metabolic communication and position yeast-derived extracellular vesicles as promising platforms for both fundamental studies of extracellular vesicle biology and emerging applications in biotechnology, metabolic engineering, and therapeutic delivery.

## Acknowledgements

We would like to acknowledge the Huck Institutes’ Metabolomics Core Facility (RRID:SCR_023864) for use of the OE 240 LC-MS and Ashley Shay and Sergei Koshkin for helpful discussions on sample preparation and analysis. We would like to acknowledge the Huck Institutes’ Cryo-EM Core Facility (RRID:SCR_024456). We would also like to acknowledge the University of Pennsylvania Penn Vet Extracellular Vesicle Core Facility (RRID:SCR_022444), and especially its director Luca Musante, PhD, for their assistance. This work was funded by the DOE Center for Advanced Bioenergy and Bioproducts Innovation (U.S. Department of Energy, Office of Science, Biological and Environmental Research Program under Award Number DE-SC0018420). Any opinions, findings, and conclusions or recommendations expressed in this publication are those of the author(s) and do not necessarily reflect the views of the U.S. Department of Energy.

## Declaration of Interests

The authors declare that they have no known competing financial interests or personal relationships that could have appeared to influence the work reported in this paper.

---

**Supplemental Figure 1.**
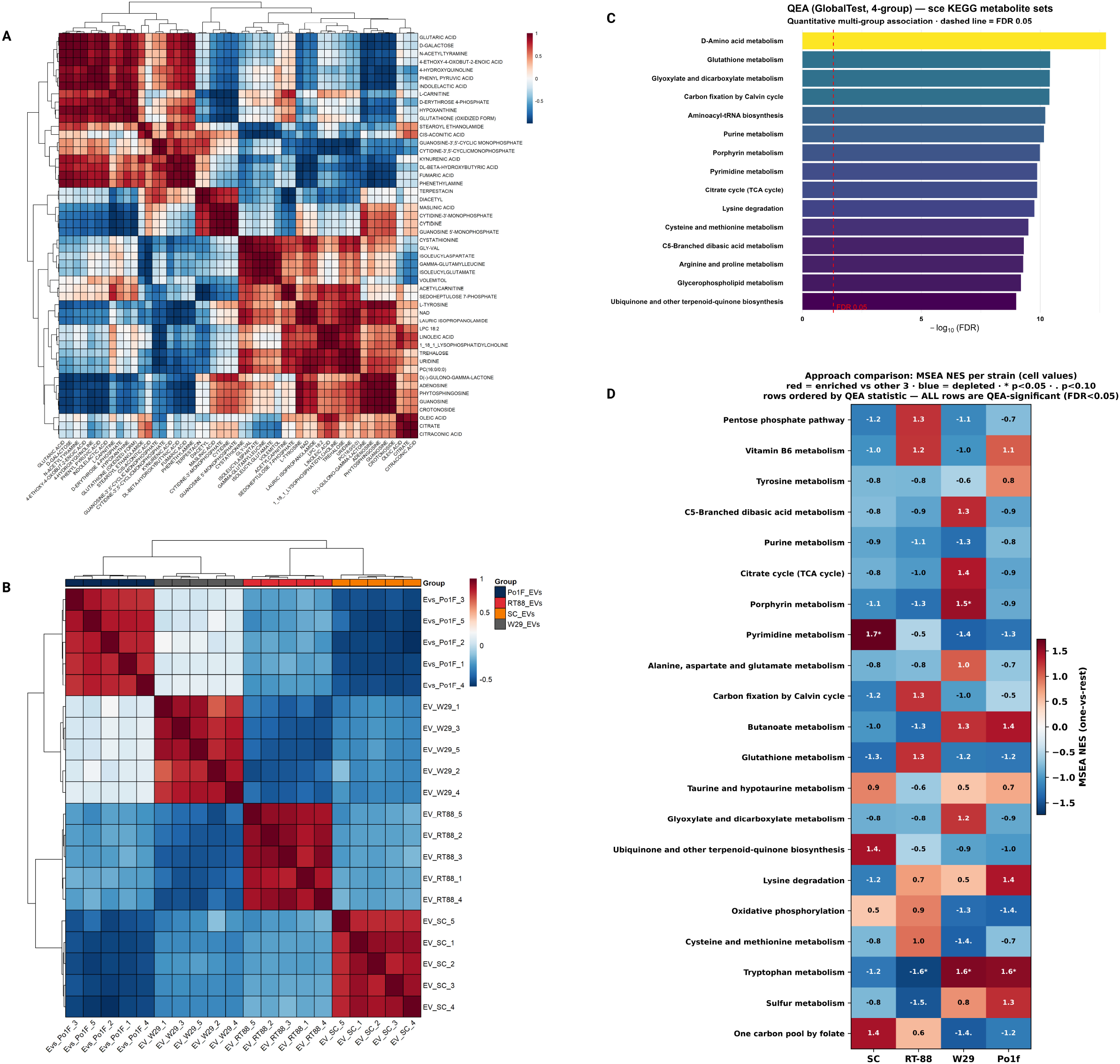

**Supplemental Figure 2.**
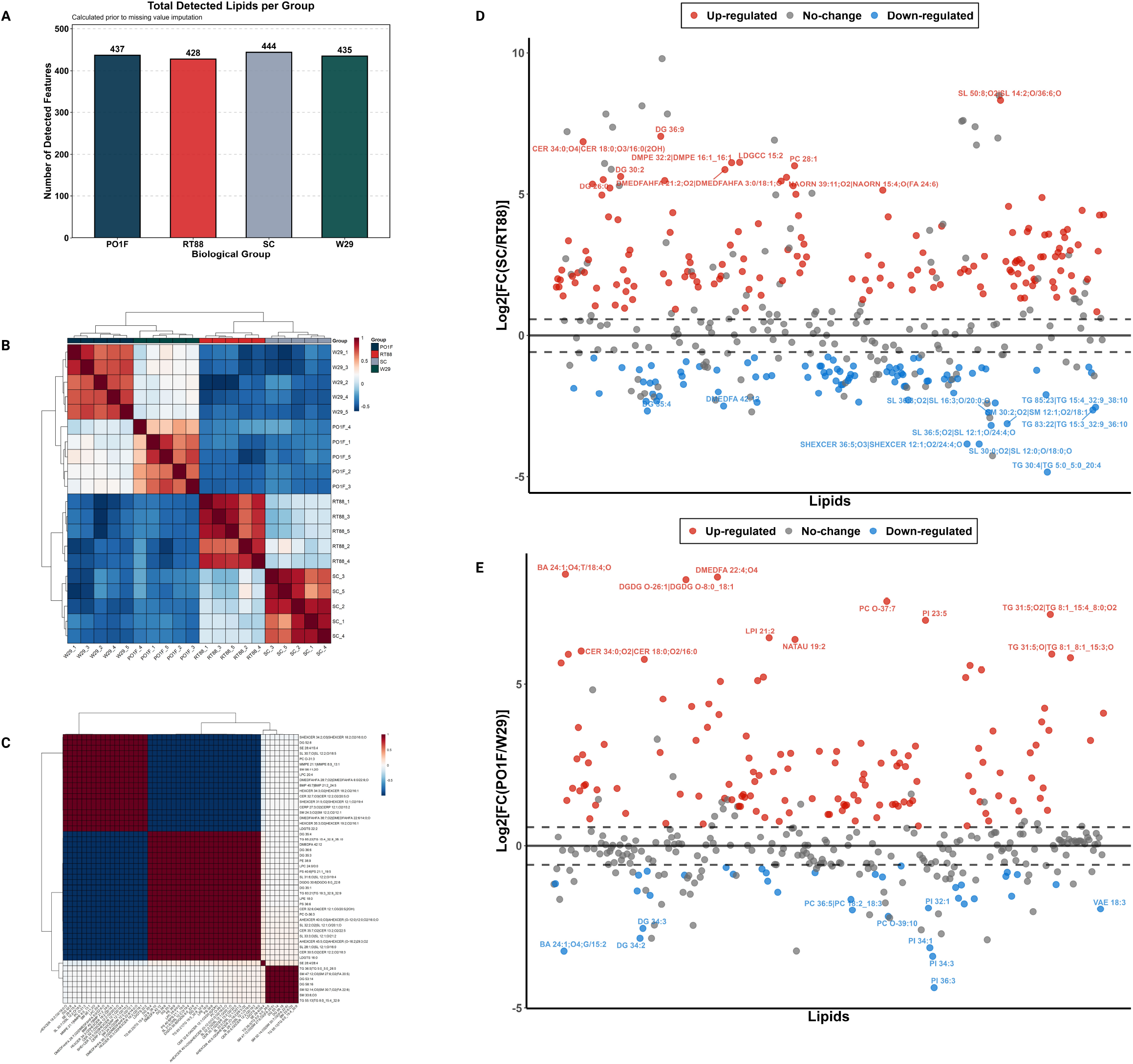

## Notes

### Competing Interest Statement

The authors have declared no competing interest.

